# Correlation Single Molecule Localization Microscopy (corrSMLM) Detects Fortunate Molecules for High Signal-to-Background Ratio and Better Localization Precision

**DOI:** 10.1101/2022.12.29.522200

**Authors:** S Aravinth, Francesca C. Zanacchi, Partha P. Mondal

## Abstract

Single-molecule localization microscopy can decipher fine details that are otherwise not possible using diffraction-limited microscopy. Often the reconstructed super-resolved image contains unwanted noise, random background and is prone to false detections. This cause spurious data that necessitates several trials, multiple experimentations, and repeated preparation of specimens. Moreover, this is not suitable for experiments that require time-lapse imaging and real-time microscopy. To overcome these limitations, we propose a technique *(corrSMLM*) that can recognize and detect fortunate molecules (molecules with long fluorescence cycles) from the recorded data. The technique uses correlation between two or more consecutive frames to extract fortunate molecules that blink for longer than the standard blinking time. Accordingly, strongly-correlated spots (single molecule signatures) are compared in consecutive frames, followed by data integration (mean centroid position and the total number of photons) and estimation of critical parameters (position and localization precision). The technique addresses two major problems that plague SMLM : (1) random noise due to false detection that contributes to strong background, and (2) poor localization precision offered by standard SMLM techniques. On the brighter side, *corrSMLM* allows only fortunate molecules contribute to the super-resolved image, thereby suppressing the background and improving localization precision by a factor of 2-4 times as compared to standard SMLM. To substantiate, corrSMLM is used for imaging fixed cell samples (Dendra2-Actin and Dendra2-Tubulin transfected NIH3T3 cells). Results show multi-fold reduction in noise and localization precision with a marked improvement in overall resolution and SBR. We anticipate *corrSMLM* to improve overall image quality and offer a better understanding of single molecule dynamics in cell biology.

## I. INTRODUCTION

In biology, molecules often assemble and arrange in a specific pattern to perform biological processes. Visualizing these patterns and understanding its arrangement is of utmost importance in fundamental biology and has ramifications ranging from cell to disease biology. In this respect, single molecule localization microscopy (SMLM) and related microscopy techniques has proven to be a useful tool for unravelling the nanoscopic world of biomolecular complexes. These powerful family of super-resolution techniques have implications in disease biology and clinical health-care [1] [2] [3].

Since the classical work of Moerner in the year 1989 and inception of super-resolution microscopy in the subsequent years with the advent of STED, SI and SMLM techniques (PALM, fPALM and STORM), super-resolution imaging has taken a giant step towards unveiling many biological processes at single molecule level [4] [5] [1] [6] [7] [8] [9] [10]. Over the years, many variants have emerged that have advanced our understanding of biological processes in a cellular environment. For example, SMLM has revealed the nanoscopic world of macromolecular complexes and its functioning in a cellular environment including, apoptopic pores [11], nuclear pore complex [12], endocytic machinery [13], cytoskeletal structure [14] and many more. Thereafter, a family of powerful techniques have emerged that has enabled detailed study of molecular complexes in live cells and organs. Recent advances in SMLM such as, POSSIBLE, STED, MINFLUX, dSTORM, SIMPLE and ROSE have shown sub-10 nm resolution [15] [18] [19] [20] [21] [22] [24]. In this regard, the integration of light sheet and super-resolution have shown great promise [16]. Specifically, the last decade has seen many variants that have immense potential to advance biology at sub-cellular level including, ground-state depletion microscopy (GSDIM) [25], super-resolution optical fluctuation imaging (SOFI) [26], points accumulation for imaging in nanoscale topography (PAINT) [27] [28], simultaneous multiplane imaging-based localization encoded (SMILE) [29] [30], individual molecule localization–selective plane illumination microscopy (IML-SPIM) [16], MINFLUX [24],POSSIBLE microscopy [15] and others [31–38]. LIVE-PAINT has demonstrated imaging a number of different peptide binding protein pairs (TRAP4-MEEVF and SYNZIP18-SYNZIP17) inside live S. cerevisiae [43]. IML-SPIM has enabled super-resolution imaging of human mammary MCF10A cell spheroids expressing H2B-PAmCherry [16]. Multicolor variant such as multicolor STORM imaging of large volumes has enabled ultrathin sectioning of ganglion cells to understand the nanoscale co-organization of AMPA receptors and their spatial correlation [39] [40] [41]. A recent technique primarily based on RESOLFT microscopy have shown 3D imaging of single molecules organization in a cell [42]. The technique has reported volumetric imaging of exosomes labelled with CD63-rsEGFP2 in live U2OS cells. Other techniques predominately based on evanescent light such as SMILE has demonstrated volume imaging capabilities in a single cell [30]. SMLM techniques are in their early stages and the emerging potential variants are expected to reveal fundamental aspects of life in the coming decade.

The last few years have seen integration of SPIM techniques with other imaging techniques for enabling studies which ere earlier thought out of reach. IML-SPIM has enabled superresolution imaging at large field-of-view (FOV). Multiphoton super-resolution microscopy. SMILE technique combines the strengths of SMLM and TIRF which facilitated 3D imaging of viral (Hemaglutinin) transfected NIH3T3 cells [29] [30]. Recently, DNA-PAINT and fluorescence lifetime PAINT (FL-PAINT) has demonstrated the capability to image multiple targets simultaneously in a HeLa cell [44]. Bessel beam STED has shown deep penetration capability that can be used for tissue / organ imaging [45]. In another application, SMLM is realized using a Bessel beam for illumination towards 3D imaging of nucleoporin Nup153 expressed in HeLa cell nucleus [46]. Recently developed MINFLUX and MINSTED techniques which is a combination of SMLM and STED has demonstrated imaging of nuclear-pore complex with high spatio-temporal resolution [49] [47] [48].

But seldom, single molecule techniques have demonstrated the ability to produce high quality images which is primarily due to strong background, sub-optimal sample preparation and large localization precision. Therefore techniques that can overcome these hurdles are highly beneficial to cell biology. Moreover, such techniques are expected to expand the reach of super-resolution technique beyond the capabilities of standard super-resolution microscopy. In this article, we propose a correlation SMLM *(corr – SMLM*) technique that uses correlation of single molecule events to identify fortunate molecules (bright molecules with large emission cycles) from consecutive frames of the recorded data. The informations (emitted photons from correlated single molecules, degree of correlation and number of correlated frames) are then integrated to determine localization precision, position of the single molecule and other parameters. Results show a multi-fold improvement in the localization precision, and high signal-to-background ratio (SBR) which facilitates imaging approaching sub-10nm regime.

## II. RESULTS

The immense potential of single molecule based superresolution microscopy can be harnessed only if the quality of reconstructed image improves substantially. Among others, localization precision and signal-to-background ratio (SBR) are key properties that are essential for high quality imaging, which becomes the basis for studying biological processes in great detail, thereby helping in deciphering the underlying mechanism. This calls for techniques / methods that are promising and does not require substantial change in system hardware. *corrSMLM* is one such general technique that can be readily integrated with most of the existing family of super-resolution microscopy techniques.

The schematic diagram of the *corrSMLM* technique / method is shown in FIg. 1. The optical system used of a standard SMLM microscopy is employed for recording several raw images. Then, the images are processed to isolate the single molecule (bright spots) which are then fitted using a 2D Gaussian function *(G(μ, σ)* = *A* exp{(*x* – *μ)*^2^/2 *σ*^2^}) where, *μ* and *σ* are the mean and standard deviation, respectively. This is followed by determining the position (centroid) and the number of photons. The centroid of extracted single molecules in each frame (say, *n)* are then compared with the single-molecule signatures in the preceding (frame #*n* – 1) and next frame (i.e, frame #*n* + 1). If the centroids are found to lie within the theoretically estimated diffraction-limited spot (*r* ∼ 1.2*λ*/2*NA*), then the respective Gaussians are correlated. Depending upon the degree of correlation, the single molecule in the consecutive frames are identified as pair and sorted as shown in the adjoining table (see, Fig. 1). Subsequently, the parameters are computed for the paired molecules. Since fortunate molecule emit for longer duration i.e, detectable in more than one consecutive frames they can be identified as a single molecule [50]. These elusive molecules has a PAR-shift towards single molecule limit, and thus have a better localization. The other benefit is the elimination of false detections (recognized as noise) which pops-in due to other reasons (detector read noise, intensity fluctuations, thermal noise etc.). Finally, the informations related to position and total number of detected photons are then used for reconstructing the single molecule map / super-resolved image.

**FIG. 1:**
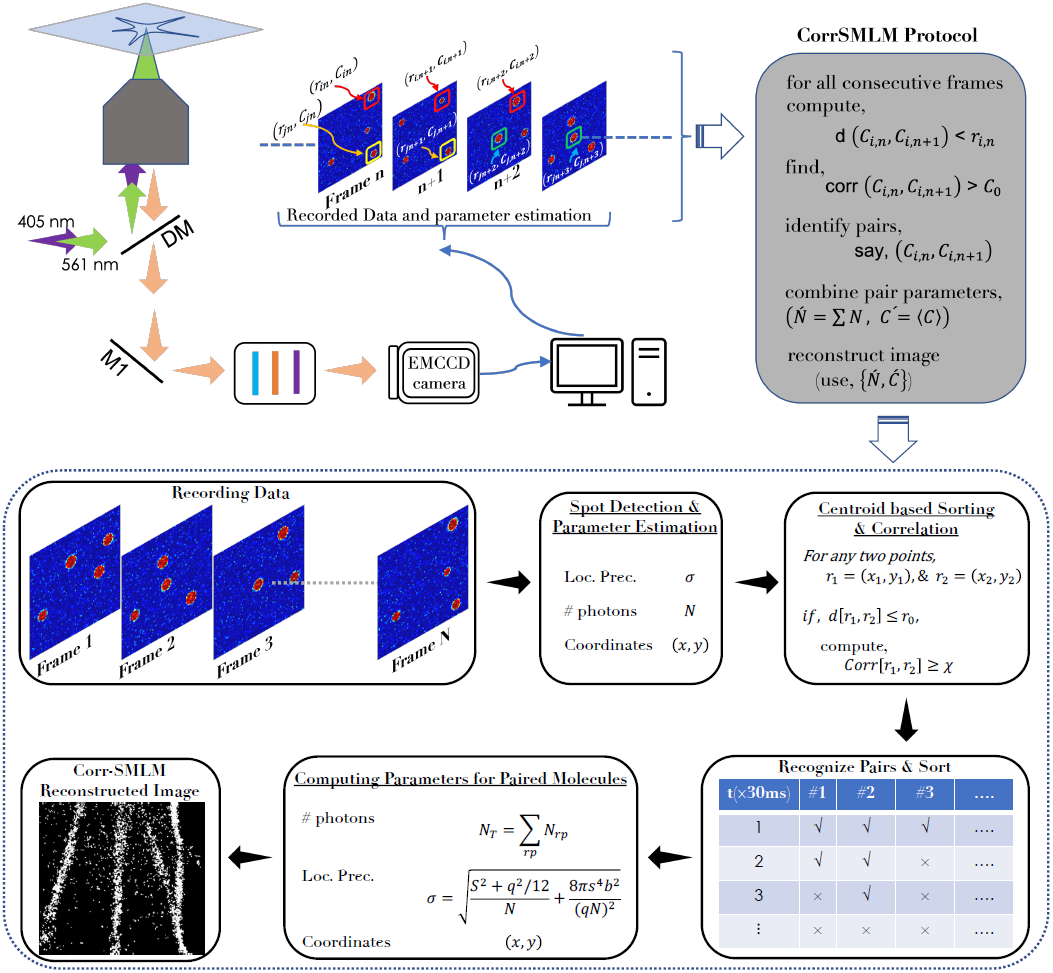
Correlation Single Molecule Localization Microscopy *(corrSMLM)*. A graphical representation of *corrSMLM* and related key steps (optical and computation). The steps involve, spot detection, recognizing and sorting fortunate molecules, correlating Gaussians (single molecule signatures) in consecutive frames, computing updated parameters (localization precision, position and number of emitted photons), and finally, reconstructing the *corrSMLM* image.

To access the potential of *corrSMLM*, two different specimens are used. We have used NIH3T3 cells and superresolved two different organelles: Actin filaments and Tubulin skeletal system in a cell. To image actin filaments, cells with a passage number of 14 were thawn and supplemented with cell medium. The cells were then cultured for two more passages to ascertain its healthy growth. Finally, the cells were prepared for transfection following standard protocol and fixed. The details related to cell culture and materials used are discussed in Ref. [3].

In the first experiment, Dendra2-Actin plasmid DNAs was used to transfect NIH3T3 cells. Subsequently, data / images are recorded by the camera at an exposure time of 33 *ms* and EM gain of 274. An estimated 49,020 images are recorded for reconstructing standard SMLM image as shown in Fig. 2. Subsequently, the images are correlated for 2, 3 and 4 successive frames. This is necessary to recognize fortunate molecules with long blinking time. The super-resolved image for *corrSMLM* is then reconstructed as shown in Fig. 2. In addition, another super-resolved image is reconstructed for 2-4 frames that combine all the single molecules detected for 2, 3, and 4 frames.

**FIG. 2:**
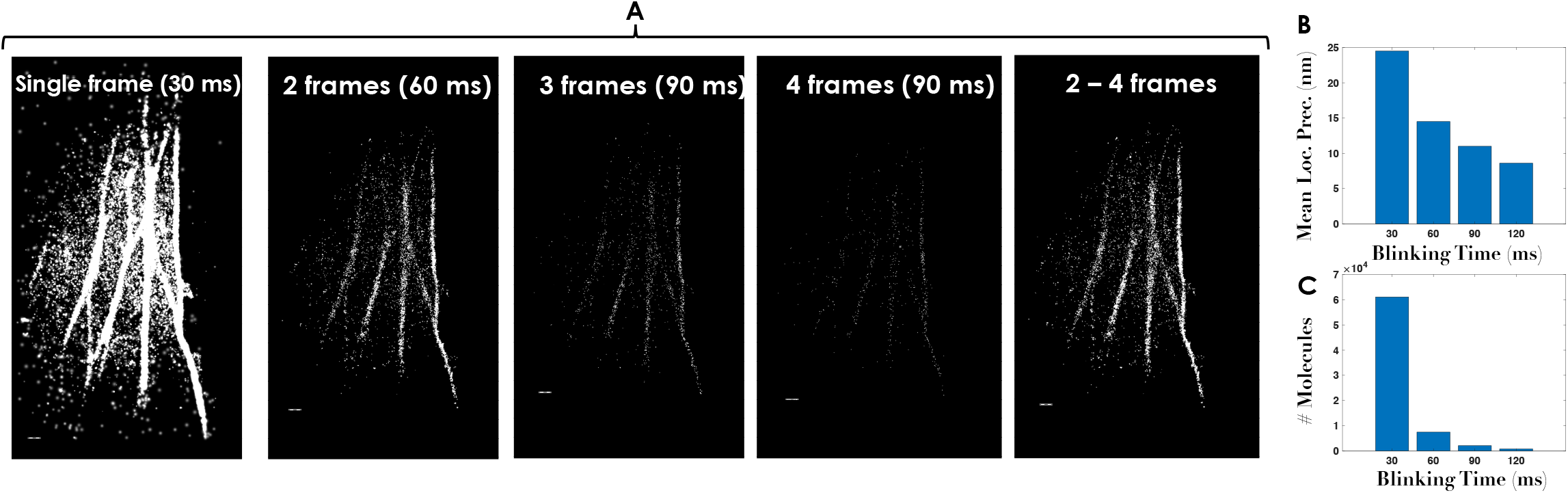
**Single molecule data collection at** 33 *Hz* **for Dendra2-Actin transfected cells, and consecutive frames are correlated to recognize fortunate molecules (single molecules that emit longer than average blinking period). Correlation was carried out for 2, 3, and 4 frames that span an exposure time of** 60, 90, 90,120 *ms*. **An additional image is reconstructed from all the single molecules collected in 2-4 consecutive frames. Corresponding statistics for mean localization precision and the number of molecules with blinking time are also shown. Scale bar** = 1 *μm*.

It becomes immediately evident that, correlated images are free from background noise, and there is a marked improvement in single molecule localization (decreased values of average localization precision as shown in Fig. 2B). This is possible due to the detection of fortunate molecules that have a better blinking period. The effect is clearly evident from the resolved actin filaments, and suppression of background noise. Another important aspect of single molecule imaging is the number of molecule that contribute to the final super-resolved image. The down-side of *corrSMLM* is probably the small number of fortunate molecules that represent the final image. Fig. 2C clearly indicates that there is a decrease in fortunate molecules that blink for longer time. This of-course reduces background and improves single molecule localization, but substantially reduces the number statistics (the number of single molecules) for acceptable super-resolved image.

The other aspect of *corrSMLM* is the degree of correlation with which two molecules can be recognized as a single-molecule blinking for longer duration (referred as pair-molecule / fortunate molecule). To ascertain, super-resolved images are reconstructed using fortunate molecules recognized by correlation factor (*χ*) of *>* 0.7, *>* 0.8 and *>* 0.9 as shown in Fig. 3A. Alongside standard SMLM image is also shown for comparison. Although the number of molecules for reconstructed image with *χ >* 0.9 is less than *χ >* 0.7 and *χ >* 0.8, key features are well preserved. The corresponding mean localization precision for all *χ*-values are also plotted in Fig. 3B. This consistently show better localization of single molecules with large blinking time and the fact that localization precision is not strongly dependent on the degree of correlation. This overall suggests that the proposed technique’s ability to accurately recognize single molecules with long blinking period.

**FIG. 3:**
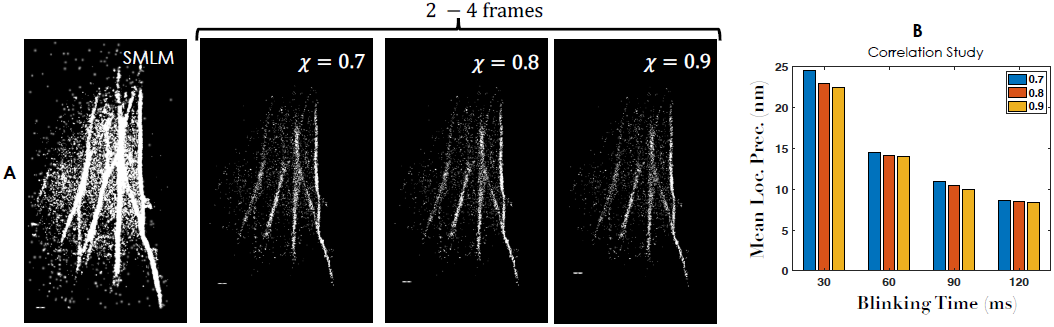
**SMLM and** *corrSMLM* **reconstructed images (integrating all single molecules in 2-4 frames) at varying correlation values**, *χ* = 0.7, 0.8, 0.9. **Alongside localization precision plot is also displayed. Scale bar** = 1 *μm*.

To strengthen our technique and additional validation, we considered another specimen where the cells are transfected with Dendra2-Tubulin plasmid DNA. A protocol similar to that of first sample (Dendra2-Actin transfected cell) is followed and the cell is fixed for imaging. Fig. 3 shows reconstructed images at varying blinking times and correlation factors, along with the localization precision. We observed a similar trend to that of first sample. It is clear that quality super-resolved image can be reconstructions with all molecules that blink for 2-4 frames. Alongside, mean localization precision with #*frames* / blinking time is also shown. It is evident that *corrSMLM* has the advantage of better number statistics, higher localization and an overall better SBR when compared to standard SMLM.

A separate experiment is carried out for comparing *corrSMLM* and standard SMLM. Fig. 5 (A,B) shows widefield fluorescence images of Dendra2-Actin and Dendra2-Tubulin transfected cells. The corresponding region-of-interest is marked (red and yellow boxes). Alongside, standard SMLM and *corrSMLM(χ* = 0.7) (combined, 2-4 frames) super-resolved images for the ROIs are also shown in Fig. 5. It is evident that, *corrSMLM* images are far superior than standard SMLM both in terms of background and resolution. To substantiate the resolving power of *corrSMLM*, we have carried out line intensity plots across actin-filaments which shows a reduction in the FWHM of actin-filaments by a factor of ∼2. In addition, we have counted the number of molecules that represent the actin filaments and background molecules in between the filaments. It is evident from the signal-to-background ratio (measured in terms of number of molecules) between two actin-fiilaments have reduced substantially when compared to standard SMLM (see, Fig. 5). This suggests overall better resolution and signal-to-background ratio for *corrSMLM*.

**FIG. 4:**
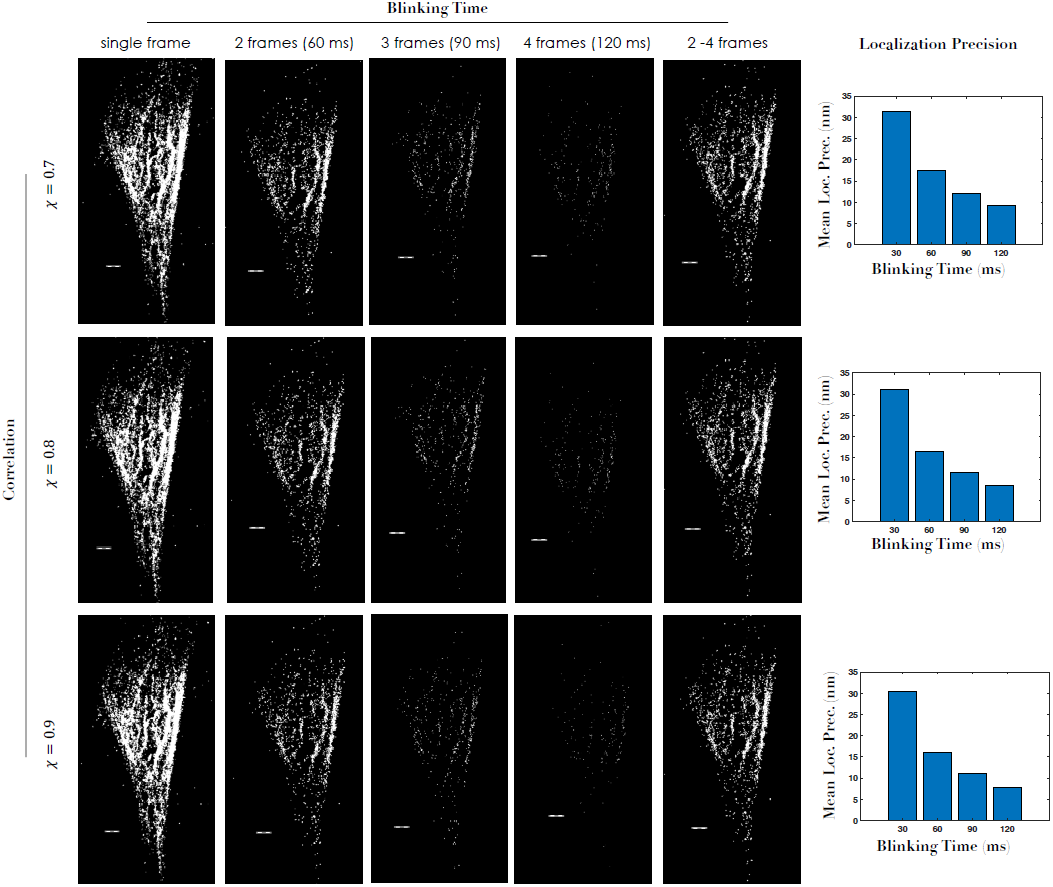
*corrSMLM* **reconstructed super-resolved images (tubulin structure labeled with Dendra2-tubulin) for 2, 3, 4 and all (2-4) frames and correlation values** (*χ* = 0.7 **to** 0.9). **Alongside localization precisions are also shown. Scale bar** = 1 *μm*.

**FIG. 5:**
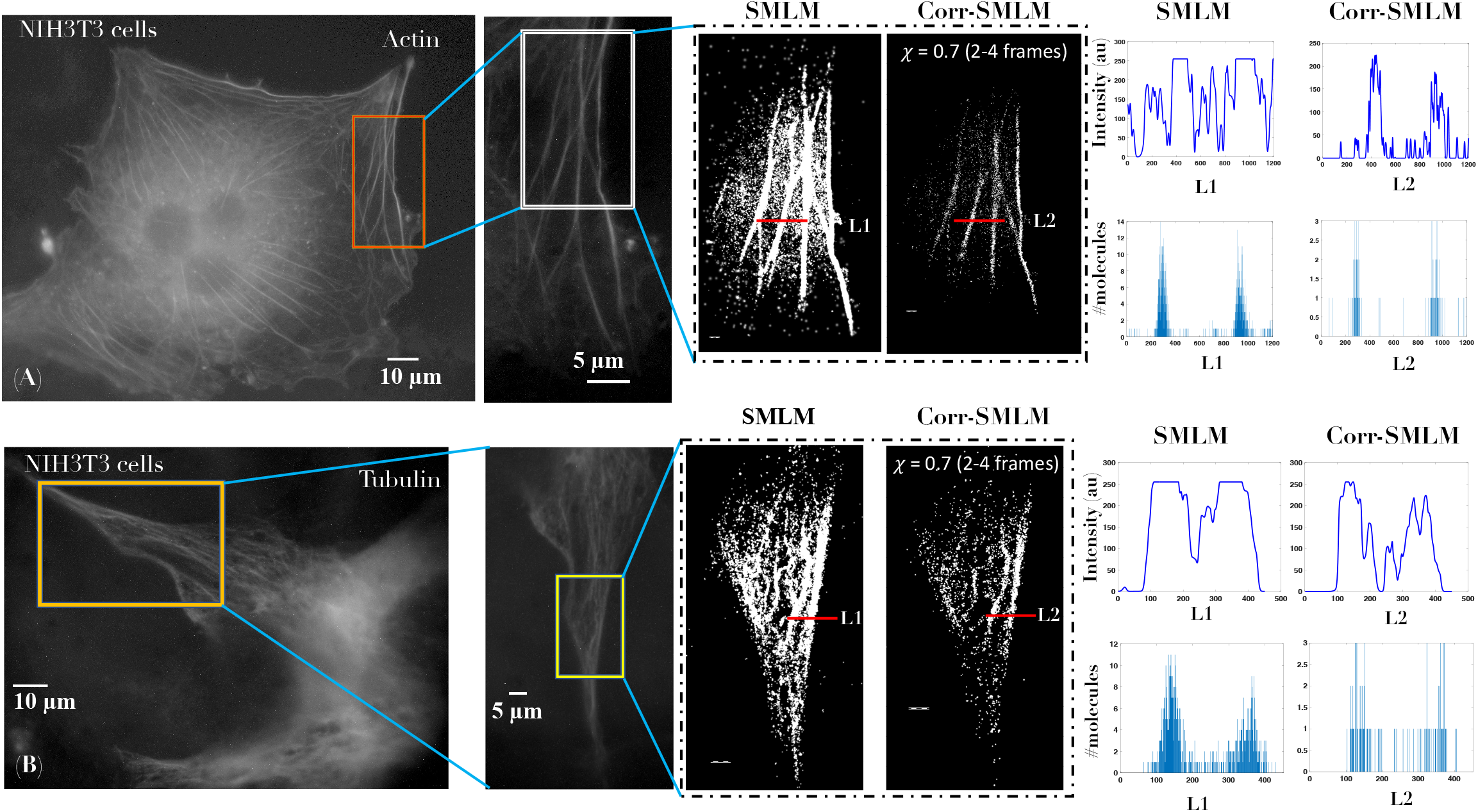
Comparison of *corrSMLM* and standard SMLM reconstructed images. The widefield fluorescence image of Dendra2-Actin and Dendra2-Tubulin transfected cells are also shown. The corresponding intensity plots along lines *L*1 and *L*2 passing through actin and tubulin structures are shown. Another plot representing the number of molecules along the lines are also shown. Scale bar = 1 *μm*.

All the findings related to *corrSMLM* are incorporated in tabular form as shown in Fig. 6. The performance of *corrSMLM* depends on few key factors such as, correlation factor, the number of correlated frames and the total number of molecules. These parameters determine the position, # photons per molecule, localization precision, and signal to background molecules. Two organelles are probed using Dendra2 photo-activable protein. It is observed that, localization of single molecules have improved by a factor of ∼2 i.e, mean localization precision is nearly-halved as compared to standard SMLM. This is closely linked to the number of emitted photons associated with the number of molecules. On the other hand, the signal-to-background ratio (measured in terms of # molecules) increased nearly 1. 5 times. Also the number of molecules necessary to represent an organelle structure (here, actin filament and tubulin network) have decreased by an order. This is predominantly due to the fact that number of molecules representing background / noise /false detections have reduced by *corrSMLM*. Overall, the benefits offers by *corrSMLM* over standard SMLM are note-worthy and useful for quality imaging.

**FIG. 6:**
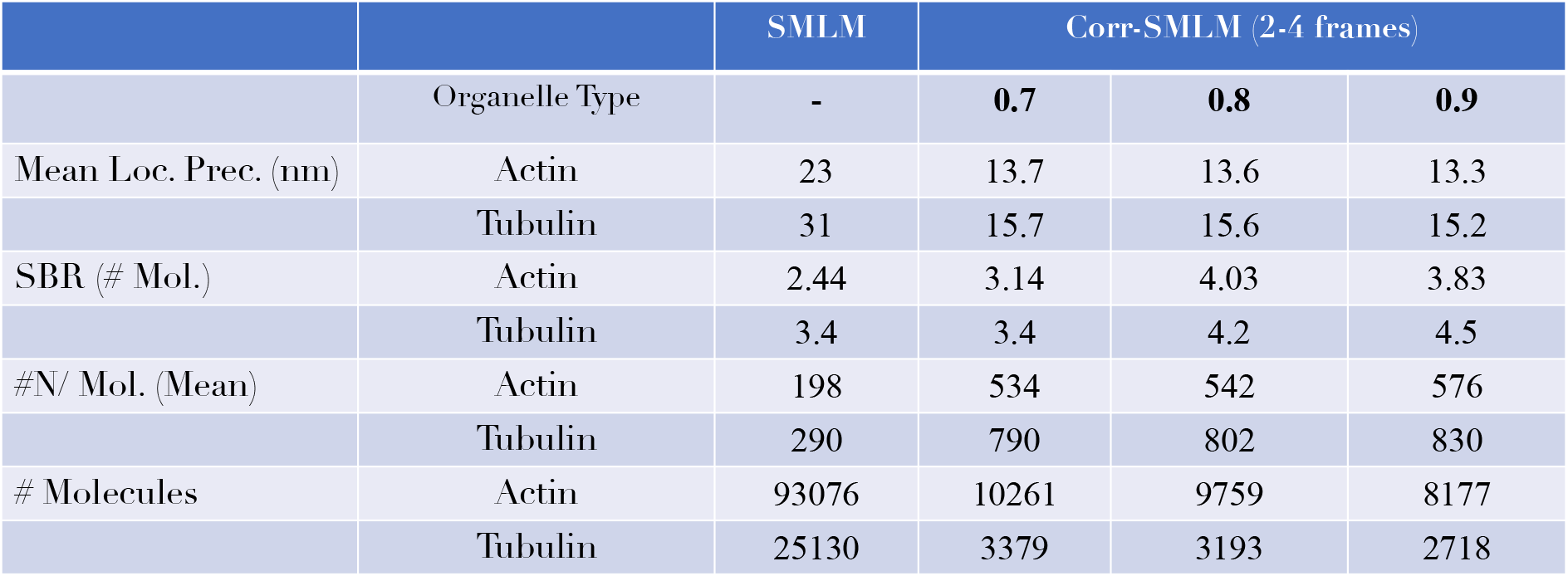
**Estimation of critical biophysical parameters (mean localization precision, signal-to-background ratio, number of emitted photons per molecule and total number of molecules) for SMLM and** *corrSMLM*.

## Discussions

A correlation-based single molecule super-resolution microscopy (*corrSMLM*) is demonstrated. The technique relies on the fact that fortunate molecules (molecules with long blinking period) stay -on for more than single data acquisition frame in SMLM, and if these elusive fluorescence events (related to fortunate molecules) are recorded, then the localization precision and signal-to-background ratio can be substantially improved. It appears that most of the genuine signal molecules are organelle specific and a fraction of these can blink for longer periods. This is unlike the background molecules or false detections that most likely get captured in single frame and are not likely to repeat in consecutive frames. Thereby, elimination of these spurious events and retaining the genuine events result in better image quality.

Single molecule experiments are performed in cellular systems following biological protocols (cell culture, transfection and fixing) to access the benefits of *corrSMLM*. We noted that plasmid DNA transfected cells express photoactivable protein (Dendra2-Actin / Dendra2-Tubulin) that are most likely located to the specific organelle (such as, actin filaments and tubulin network). Any other events (representing single molecules) are most likely due to noise (intensity fluctuation or false detections), background and detector inefficiencies (read / thermal noise). When two or more consecutive frames are correlated these noises are filtered out, thereby retaining only the genuine molecules. Hence, the resultant reconstructed images are free from background. This is quite evident from Fig. 2 and Fig. 4. Another important aspect of *corrSMLM* is the correlation factor (taken between 0.7 to 0.9) for which the corresponding reconstructed super-resolved image appears to represent the target organelle (see, Fig. 3). This is encouraging since this emboldens the fact that there is a strong correlation of few single molecule events in successive recorded frames.

Finally, the proposed *corrSMLM* technique is compared with the standard SMLM. Analysis show improvement in resolution and signal to background molecules when investigated using line intensity plots across actin filament and tubulin structures (see, Fig. 5). This directly impacts resolution and overall image quality. Specific analysis of key aspects suggests the benefits of *corrSMLM* over SMLM in terms of mean localization precision, signal-to-background ratio (#Mol) among others. With all the benefits and non-requirement of additional hardware, *corrSMLM* is expected to advance localization based super-resolution microscopy.

## Contributions

All authors contribute equally to this work.

